# Individual-based multiple-unit dissimilarity: novel indices and null model for assessing temporal variability in community composition

**DOI:** 10.1101/2021.01.17.427031

**Authors:** Ryosuke Nakadai

## Abstract

Beta-diversity was originally defined spatially, i.e., as variation in community composition among sites in a region. However, the concept of beta-diversity has since been expanded to temporal contexts. This is referred to as “temporal beta-diversity”, and most approaches are simply an extension of spatial beta-diversity. The persistence and turnover of individuals over time is a unique feature of temporal beta-diversity. Nakadai (2020) introduced the “individual-based beta-diversity” concept, and provided novel indices to evaluate individual turnover and compositional shift by comparing individual turnover between two periods at a given site. However, the proposed individual-based indices are applicable only to pairwise dissimilarity, not to multiple-temporal (or more generally, multiple-unit) dissimilarity. Here, individual-based beta-diversity indices are extended to multiple-unit cases. In addition, a novel type of random permutation criterion related to these multiple-unit indices for detecting patterns of individual persistence is introduced in the present study. To demonstrate the usage the properties of these indices compared to average pairwise measures, I applied them to a dataset for a permanent 50-ha forest dynamics plot on Barro Colorado Island in Panama. Information regarding “individuals” is generally missing from community ecology and biodiversity studies of temporal dynamics. In this context, the methods proposed here are expected to be useful for addressing a wide range of research questions regarding temporal changes in biodiversity, especially studies using traditional individual-tracked forest monitoring data.

## Introduction

The concept of beta-diversity was introduced by Whittaker (1960, 1972) to define the variation in community composition among sites in a region. Several other beta-diversity indices have subsequently been proposed (reviewed by Koleff et al. 2003; Anderson et al. 2011; Jost et al. 2011; Legendre and De Cáceres 2013). Recent studies have focused on temporal changes in community composition at both single sites and multiple sites surveyed repeatedly over time (Magurran 2011; Legendre and Condit 2019). Temporal changes in community composition are referred to as “temporal beta-diversity” (Hatosy et al. 2013; Legendre and Gauthier 2014; Shimadzu et al. 2015), which is simply an extension of spatial beta-diversity. Only a few studies have focused on methodological developments of temporal beta-diversity (Shimadzu et al. 2015; Legendre 2019; Nakadai 2020). The persistence and turnover of individuals over time is a key feature of temporal beta-diversity (Magurran et al. 2019; Nakadai, 2020). The speed and frequency of compositional change over time is associated with the speed of individual turnover, and must be considered in community comparisons because even randomly high individual turnover can result in high temporal beta-diversity (Nakadai 2020).

To address this issue, Nakadai (2020) proposed the “individual-based beta-diversity” concept, and provided novel indices to evaluate individual turnover and compositional shift by comparing individual turnover between two periods at a given site. Specifically, I developed an individual-based temporal beta-diversity approach for the Bray–Curtis dissimilarity index (i.e., an abundance-based temporal beta-diversity index) (Nakadai 2020). Over time, some individuals replace others, and therefore ecological communities are dynamic and vary to some degree both spatially and temporally (Mori et al. 2018; Tatsumi et al. 2019). In addition, some individuals of long-lived species (e.g., trees and vertebrates) persist over the long term, whereas individuals of short-lived species are replaced with conspecifics or individuals of other species (Nakadai 2020).

The average values of pairwise dissimilarity have been widely used to evaluate compositional heterogeneity among multiple sites but cannot accurately reflect the overall compositional heterogeneity within a pool of more than two sites (Baselga 2013a). Diserud and Ødegaard (2007) first proposed a multiple-site (or more generally, a multiple-unit) similarity measure, which has conceptual and methodological advantages over conventional approaches based on average pairwise similarities (Koleff and Gaston 2002; Gaston et al. 2007; Baselga et al. 2007). Many dissimilarity indices have been extended to multiple-unit indices (Baselga 2013b, 2017). Multiple-unit dissimilarity indices can take into account components shared among more than three units (i.e., sites or temporal units) (Baselga et al. 2007). The pairwise individual-based indices proposed by Nakadai (2020) may be limited by the typical problems associated with the application of multiple-unit datasets. In particular, even if the numbers of mortal and recruited individuals are same, the presence of long-lived individuals could result in underestimates of individual turnover when the average values of the pairwise indices were used. Extending individual-based indices to multiple-temporal datasets may allow accurate assessment of the temporal variability of community composition based on individual turnover.

A null model is a pattern-generating model based on randomization of ecological data and is designed to produce a pattern that would be expected in the absence of a particular ecological mechanism (Gotelli and Graves 1996; Gotelli and McGill 2006). Null models for temporal patterns in ecological communities have been evaluated previously (e.g., cyclic shift permutations; Hallett et al. 2014, 2016). Such models use a matrix of the abundance, presence or absence, or biomass of species over multiple temporal units (e.g., years) as an input. In each realization of the algorithm, the time series of every species is shifted forward by a random number of units, independently of other species (Kalyuzhny 2020). However, no null model approach using individual randomization has been developed for community dynamics, and no research has been conducted on the patterns of individual persistence using either beta-diversity indices or null models.

Therefore, the purpose of this paper was to develop new multiple-unit indices that can quantify compositional variability across time according to the speed of individual turnover, as well as novel random permutation criteria to evaluate the deviation in individual persistence from the perspective of ecological drift. In the present study, I extended individual-based beta-diversity indices to multiple units after briefly reviewing the history of pairwise and multiple-unit indices and also introduced a newly developed null model. Then, I applied these indices to data from a 50-h forest dynamics plot on Barro Colorado Island (BCI) in Panama (Condit et al. 2019), as a case study to demonstrate the usage and properties of indices compared to average pairwise measures. Using both novel indices and a null model provides opportunities to answer the following questions pertaining to the BCI plot. Are differences in the degree of compositional variability explained only by differences in the degree of individual turnover? Does the degree of deviation in individual persistence from each expected value of ecological drift: lead to differences in compositional variability among forest plots?

## Materials and methods

### Conventional incidence-based and abundance-based indices and novel individual-based indices

Two types of dissimilarity indices have been proposed in previous studies: incidence-based and abundance-based dissimilarity indices (Baselga 2013b; Nakadai, 2020). Specifically, the widely used Bray–Curtis dissimilarity index is an abundance-based extension of the Sørensen index (Legendre and Legendre 2012; Baselga 2013b). The Bray–Curtis dissimilarity index was originally proposed as a percentage difference index by Odum (1950), yet it is commonly misattributed to Bray and Curtis (1957). The incidence-based and abundance-based methods target the number of species and the number of individuals, respectively. Conventional beta-diversity indices are based on intersection (*A, a*) and relative complement (*B, b* and *C, c*) components. For example, the Sørensen dissimilarity index (*d_sor_*; Sørensen 1948) is formulated as follows:

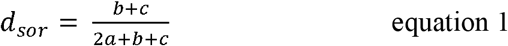

where *a* is the number of species common to both units (i.e., sites or temporal units), *b* is the number of species that occur in the first but not the second unit, and *c* is the number of species that occur in the second but not the first unit. Similarity, the Bray–Curtis dissimilarity index (Bray and Curtis 1957) is formulated as follows:

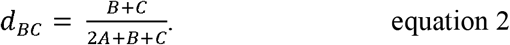

The abundance-based components are as follows:

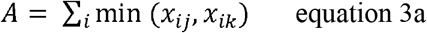

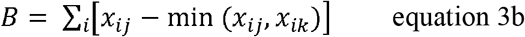

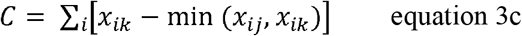

where *x_ij_* is the abundance of species *i* at unit (i.e., site or time) *j*, and *x_ik_* is the abundance of species *i* at unit (i.e., site or time) *k*. Therefore, *A* is the number of individuals of each species common to both units *j* and *k*, whereas *B* and *C* are the numbers of individuals unique to units *j* and *k*, respectively.

Therefore, it should be possible to distinguish the community dynamics component of persistence (*P*) from those that change over time, i.e., mortality and recruitment (*M* and *R*, respectively). *M* refers to individuals occurring at time 1 (abbreviated T1) that die before time 2 (abbreviated T2), thereby representing the death of individuals during the time between T1 and T2. *R* identifies individuals that were not alive at T1 (or were not yet in the size class counted, e.g., diameter breast height ≥ 1 cm) but were subsequently alive and counted at T2, thereby representing *R* between T1 and T2. Here, the term “recruitment (*R*)” is used because I have tree communities in mind. “Colonization” in the sense of Tatsumi et al. (2021) could be employed when the approach introduced in this study is applied to studies on taxa other than trees (e.g., bird ringing data). Component *P* refers to the persistence of individuals from T1 to T2. Although *P, M*, and *R* relate to components *A, B*, and *C*, respectively, here I have used different characters to emphasize the processes represented by each component. The respective abundances were calculated as follows:

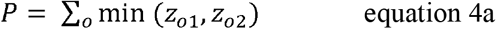

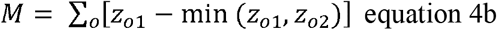

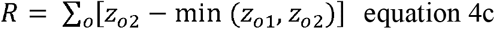

where *z_o1_* is individual *o* at time T1, and *z_o2_* is individual *o* at time T2. Both *z_o1_* and *z_o2_* take a value only of 1 or 0 (presence and absence, respectively). Therefore, *P* is the total number of individuals present at both T1 and T2, whereas *M* and *R* are the numbers of individuals unique to T1 and T2, respectively (Nakadai 2020).

T1 and T2 can be generalized as unit *j* (T*j*) and unit *k* (T*k*), and generalized versions of *P, M*, and *R* for units *j* and *k* are simply formulated as follows:

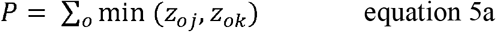

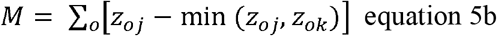

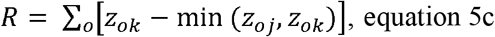

which are used for the main analysis of this study.

This formulation of the individual-based temporal beta-diversity index for Bray–Curtis (*d_MR_*) dissimilarity can be expressed as follows:

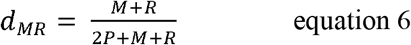

In some cases, there is turnover of different individuals belonging to the same species even when the community composition is stable over time, which contributes to a dynamic compositional equilibrium across time periods. Therefore, the components that change over time (*M* and *R*) are re-organized into two further components: compositional equilibrium (*E*) and shift (*B* and *C*), respectively (Nakadai 2020; Fig. 1). Component *M* can be partitioned into lost individuals that contribute to compositional change in a community (*B*) and lost individuals replaced by conspecifics, contributing substantially to equilibrium in community composition (*E_loss_*) (Fig. 1). Similarly, component *R* can be partitioned into gained individuals that contribute to compositional change in a community (*C*) and gained individuals replacing conspecifics, thereby contributing substantially to equilibrium in community composition (*E_gain_*) (Fig. 1). By definition, the numbers of *E_loss_* and *E_gain_* individuals are identical; hence, I replaced both *E_loss_* and *E_gain_* with the coefficient *E*, i.e., the component that contributes to dynamic compositional equilibrium (Nakadai 2020). Furthermore, components *M* and *R* can also be analyzed in the same manner as community data (Nakadai 2020). Therefore, the Bray–Curtis dissimilarity between communities can be calculated based on *M* and *R* as follows:

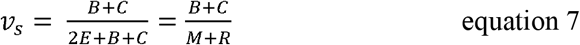

*where v_s_* indicates the speed of compositional shifts in a community relative to the total speed of individual turnover associated with *M* and *R*. If *v_s_* is 0, then individual turnover would not contribute to compositional shifts in a community; i.e., *M* and *R* are identical, and an equilibrium state exists. If *v_s_* is 1, all individual turnovers contribute to compositional shifts, and *M* and *R* are completely different. It is also possible to calculate *v_s_* as *d_BC_* divided by *d_MR_*, as follows:

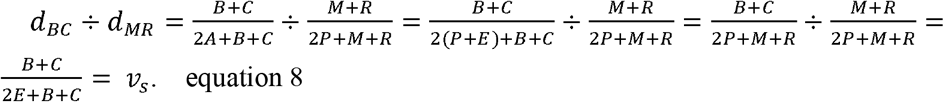

**Figure 1.**
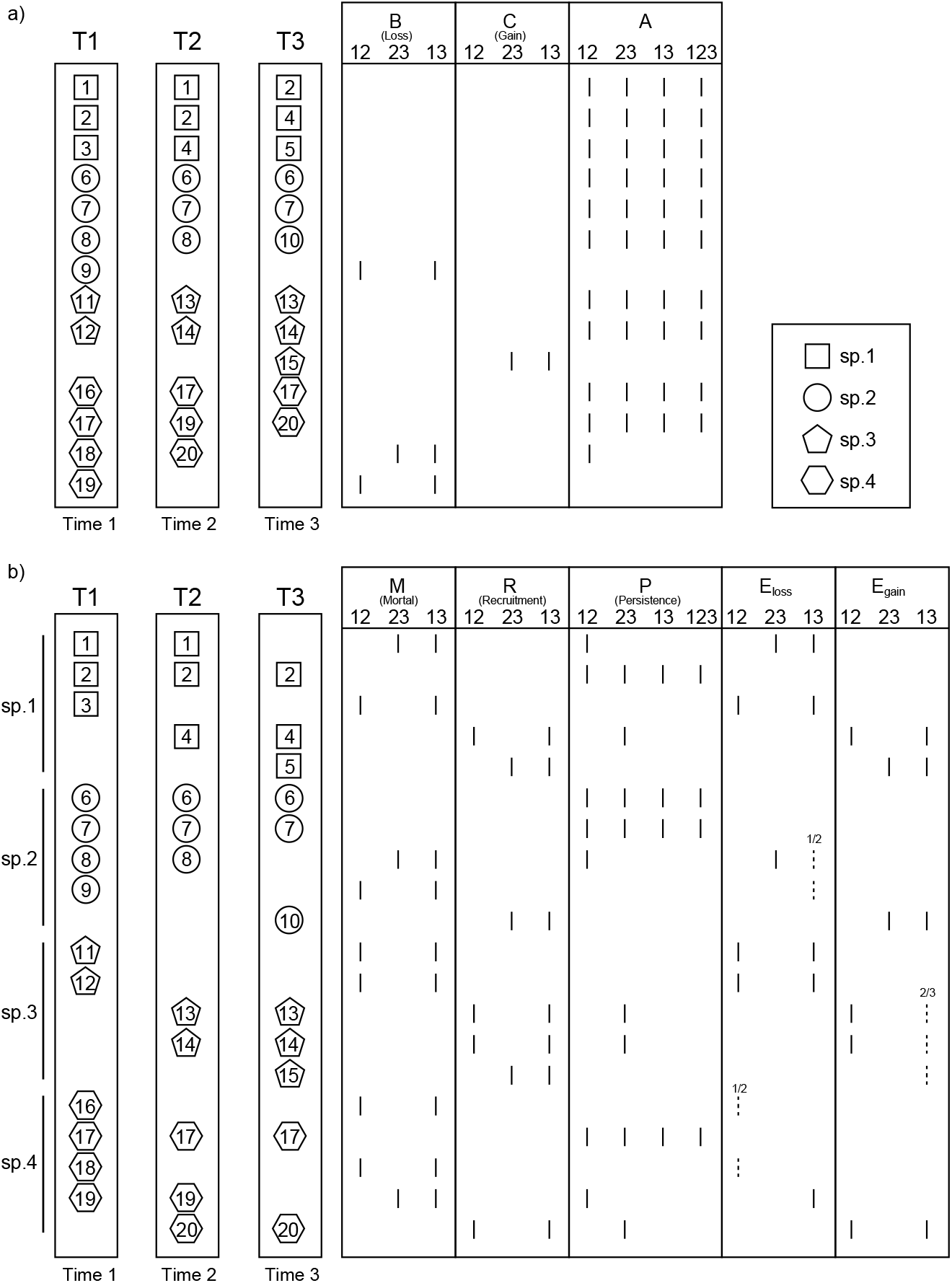
Comparison of species composition and individuals (20 individuals of four species total) among temporal units, showing the components (*A, B, C, M, R, P, E_loss_, and E_gain_*) used for assessing compositional variability over time. The squares, circles, pentagons, and hexagons indicate species 1, 2, 3, and 4, respectively. The numbers under each element (e.g., 12) represent the corresponding time step. For components *A* and *P*, the number 123 is also shown, indicating the shared species and individuals across the three time steps, respectively. Detailed explanations of each component are provided in Table 1. The dashed lines with numbers (i.e., a denominator and a numerator) indicate that the numerator chosen from among candidates of the same species (i.e., the number in the denominator) will be *E_loss_* or *E_gain_* because the components of *E_loss_ and E_gain_* do not require individual identity information.

**Table 1.**
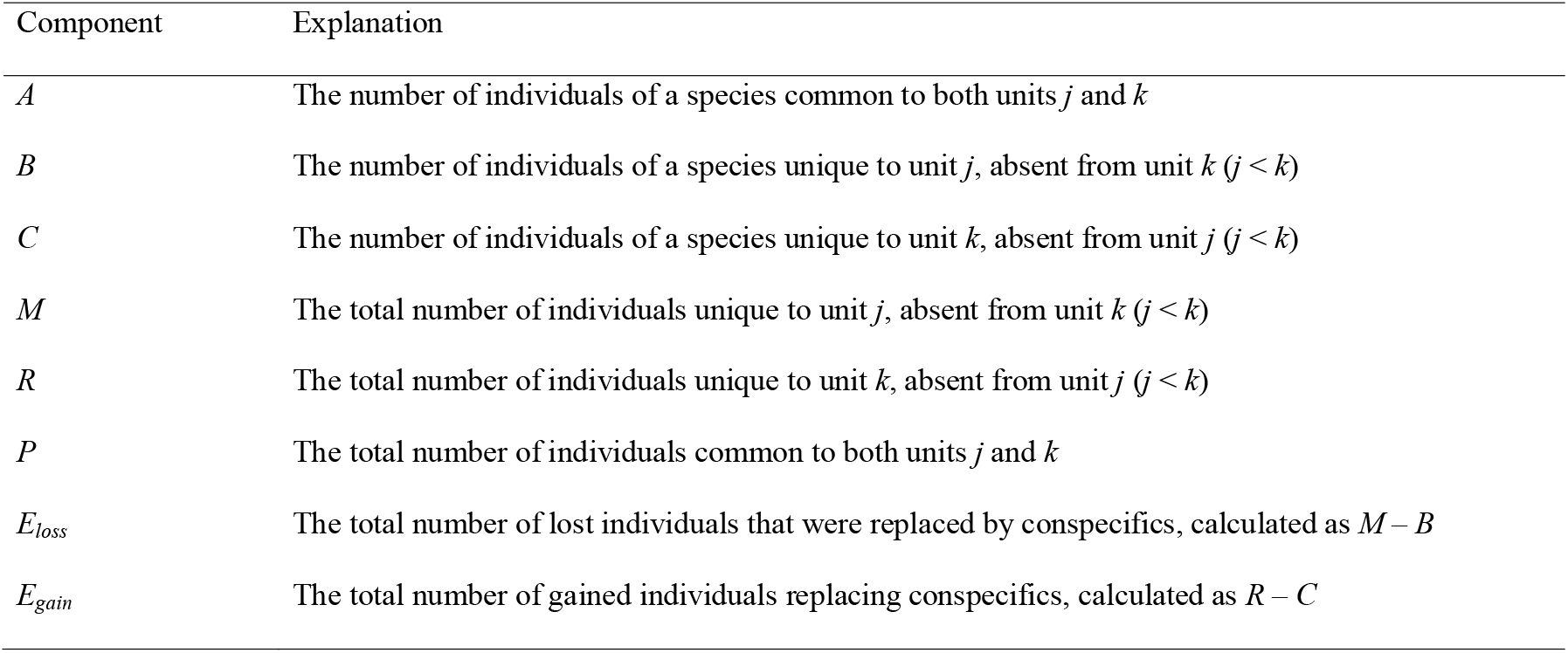
Explanations of the components used in the conventional and novel indices of beta diversity. A visualization of those components is provided in Fig. 1.

**Table 2.**
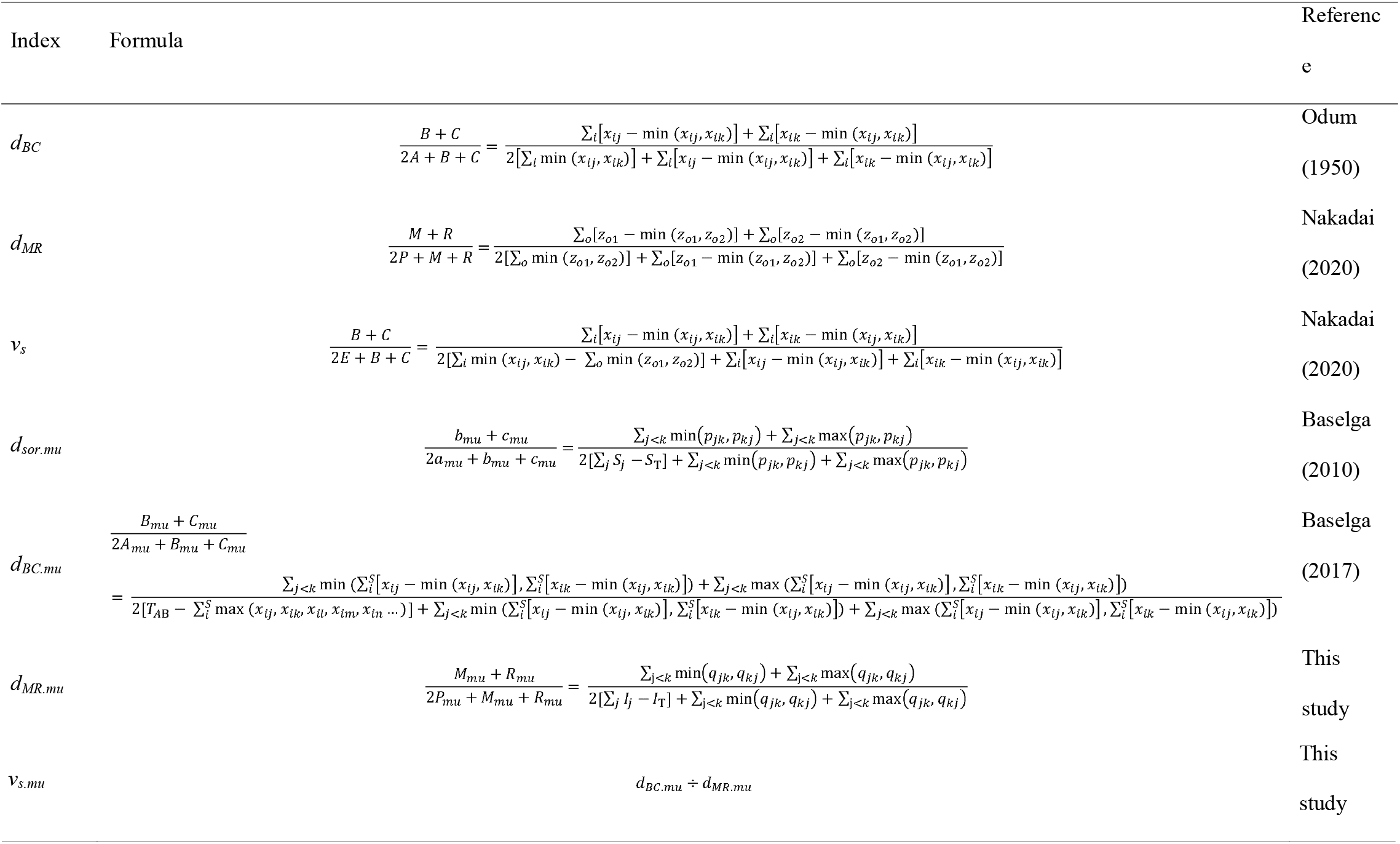
Summary of indices, formulae, and references citing both the indices developed in previous studies and the novel indices proposed in this study.

### Conventional multiple-unit beta-diversity indices

Multiple-site dissimilarity indices were first proposed by Diserud and Ødegaard (2007), which made it possible to consider species information shared among more than three units using the inclusion–exclusion principle (Erickson, 1996). Since then, various multiple-unit beta-diversity indices have been proposed (Baselga et al. 2007; Baselga, 2013b), including several new multiple-unit indices based on pairwise dissimilarity indices (Baselga et al. 2007; Baselga 2010, 2012, 2017). Specifically, these multiple-unit dissimilarity indices share the *A (a), B (b)*, and *C (c)* components of indices based on abundance (incidence). *a*_mu_, *b_mu_*, and *c_mu_* correspond to *a, b*, and *c*, respectively, of incidence-based pairwise dissimilarity indices.

Although both *b_mu_* and *c_mu_* are simply sums of all components divided into two parts in the pairwise indices, *a_mu_* is based on the inclusion–exclusion principle described above and is difficult to understand intuitively. For example, in the case of three units, the component *a*_mu_ can be calculated as follows (Baselga et al. 2007; Baselga, 2010):

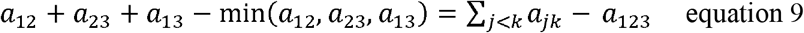

where *a_jk_* is the number of species common to both units. Considering a general case with *n* units, the component *a_mu_* can be formulated as follows:

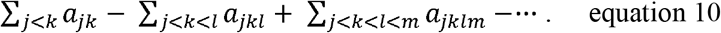

Equation 9 can be converted based on the inclusion–exclusion principle, and the resulting components are as follows:

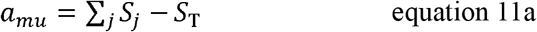

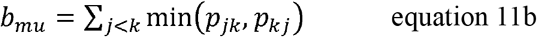

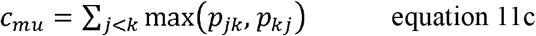

where *S_j_* is the total number of species at site *j, S_T_* is the total number of species at all sites considered together, and *p_jk_* and *p_kj_* are the numbers of species exclusive to units *j* and *k*, respectively, when compared pairwise (Baselga 2010). Specifically, *p_jk_* is the number of individuals included in unit *j* but not in unit k, and *vice versa*.

Similar to the case of *a_mu_* described above, the component *A_mu_* can be calculated for *n* units as follows (Baselga 2010):

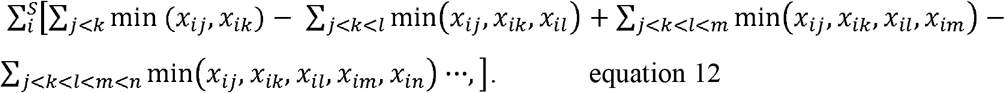

Applying the inclusion–exclusion principle allows the *A_mu_* component to be simplified. *A_mu_*, *B_mu_*, and *C_mu_* correspond to *A, B*, and *C*, respectively, from abundance-based pairwise dissimilarity indices:

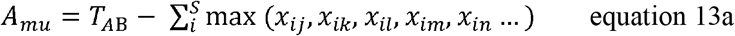

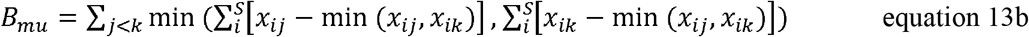

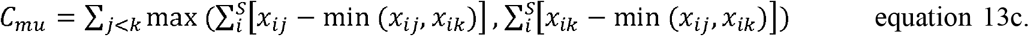

where *T_AB_* is the total abundance in the data set (details see Baselga, 2017).

Multiple-unit indices for Sørensen and Bray–Curtis are formulated as follows (Baselga 2010, 2013b),

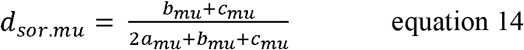

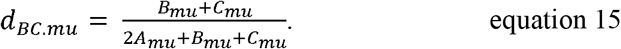

### Novel individual-based multiple-unit beta-diversity indices

In this study, I developed individual-based components and novel indices for multiple-unit data sets. In accordance with existing methods for the calculation of multiple-unit dissimilarity, the persistence (*P_mu_*) component can be calculated for three time steps as follows:

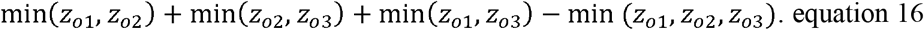

Considering a general case with *n* units, the component *P*_mu_ can be formulated as

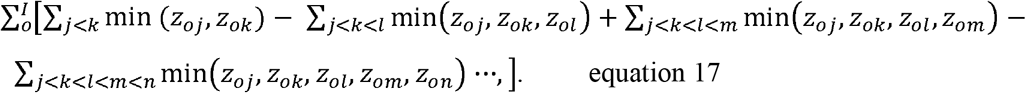

Using the inclusion–exclusion principle, *P_mu_* can be simplified and the components *P_mu_, M_mu_* and *R_mu_* reformulated as follows:

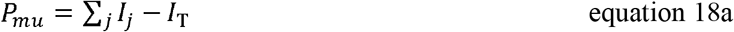

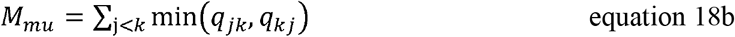

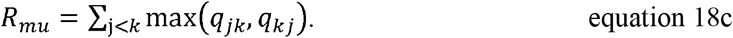

where *I_j_* is the total number of individuals in a unit *j, I_T_* is the total number of individuals in all units considered together, and *q_jk_* and *q_kj_* are the numbers of individuals exclusive to units *j* and *k*, respectively, when compared pairwise.

The multiple-unit individual-based index is formulated as follows:

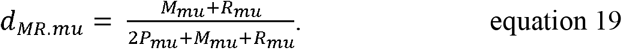

Furthermore, as an extension of equation 8, I also introduced *v*_s.mu_, which indicates the compositional variability in a community relative to the individual turnover, as follows:

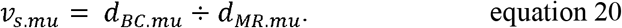

### Novel randomization criteria to construct a null model of individual-tracked monitoring data

Here, I introduce a novel method for constructing a null model to evaluate the degree of individual persistence over time. Figure 2 is a conceptual diagram outlining the basis of the new null model. Cases (i) and (ii) shown in Fig. 2 illustrate the patterns of individual persistence of two individuals. The degree of individual persistence differs between the two cases, even if the numbers of individuals involved in recruitment and mortality were equal across the time steps. In both cases, one individual was recruited, and one individual died. Specifically, in case (i), the individual *z_1_* persists over time, whereas individual *z_2_* was recruited between T1 and T2 and died between T2 and T3. On the other hand, in case (ii) described below, individual *z_1_* died between T2 and T3, and *z_2_* was recruited between T1 and T2 and persisted, at least until the end of observation at T4. Such differences in the patterns of individual persistence lead to differences in the value of the newly developed individual-based temporal multiple-unit dissimilarity index (*d_MR.mu_*), even when the numbers of individuals undergoing mortality and recruitment are equal. The component *P*_mu_ can account for differences in the pattern of individual persistence. Specifically, *P*_mu_ decreases, and the index *d_MR.mu_* increases, when the number of long-lived individuals (i.e., more than three time steps) increases, and *vice versa*. The null model approach introduced here uses those properties of the component and the index.

**Figure 2.**
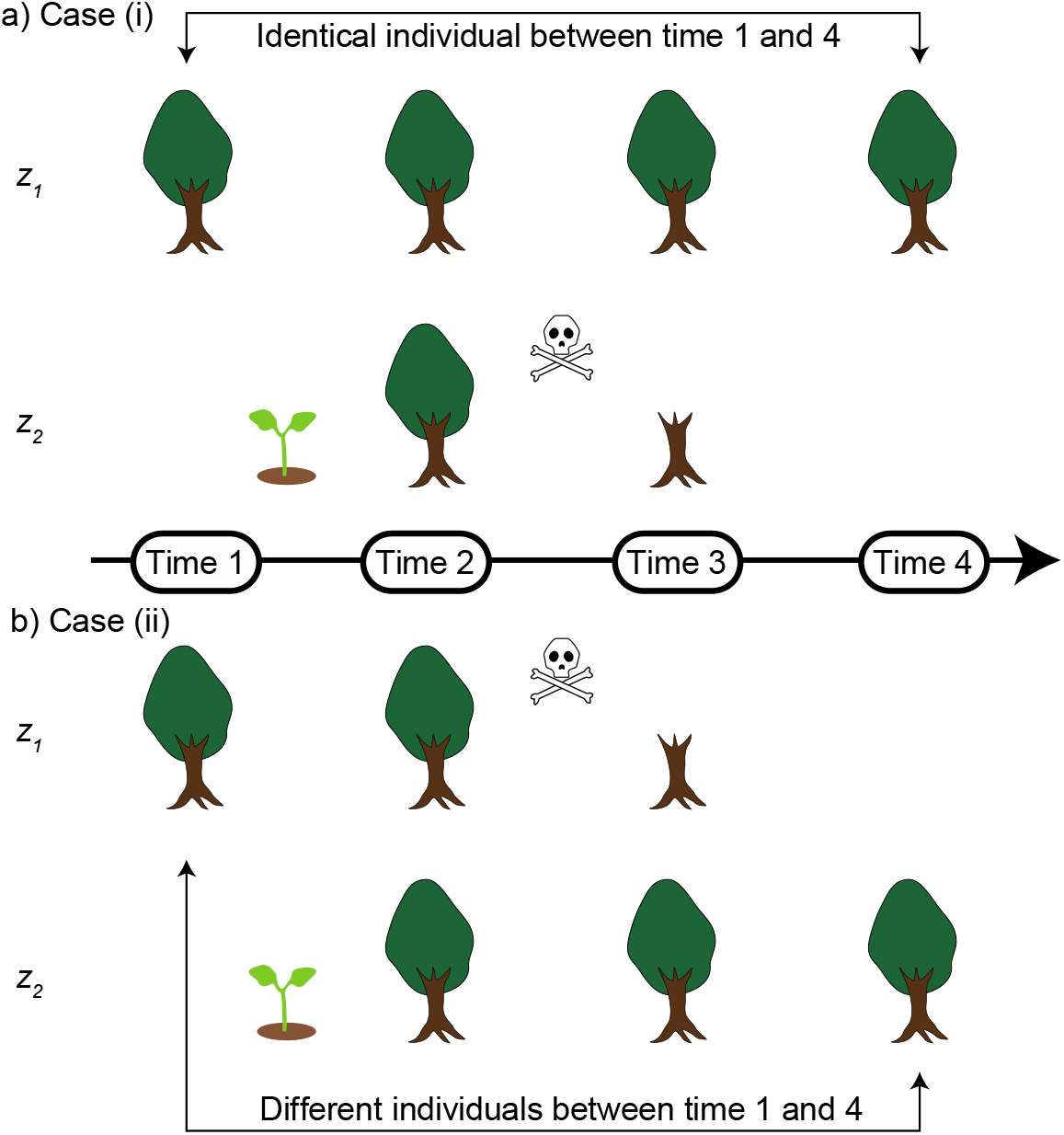
Conceptual diagram showing the basis of the new null model. Both a) case (i) and b) case (ii) showed patterns of individual persistence for two individuals. The pattern of individual persistence differed between the two cases, even if the numbers of individuals affected by recruitment and mortality were the same at each time step. Here, in both cases, one individual was recruited, and one individual died. Specifically, in case (i), individual *z_1_* persisted over time, but individual *z_2_* was recruited between times 1 and 2 and died between times 2 and 3. On the other hand, in case (ii), individual *z_1_* died between times 2 and 3, and individual *z_2_* was recruited between times 1 and 2 and continued to persist at least until time 4. The average value of the individual-based temporal beta-diversity index 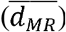, which indicates the degree of individual turnover, may be the same in both cases, but the newly developed multiple-unit dissimilarity index will have different values (*d_MR.mu_*) because the component *P*_mu_ can account for differences in the pattern of individual persistence. Specifically, *P*_mu_ decreases, and the index *d_MR.mu_* increases, when the number of long-lived individuals (i.e., those persisting for more than three time steps) increases. The sprout and the skull and crossbones symbols indicate recruitment and mortality, respectively.

In the novel null model, the simulations use the original numbers of recruited and mortal individuals, but mortal individuals are randomly selected at each time step. The simulations were repeated (e.g., 100 times) for each plot. The index *d_MR.mu_* can be calculated from each repeated simulation. By comparing the observed and simulated values of the index *d_MR.mu_*, the standardized effect size of the degree of deviation in individual persistence from the null model (*P_ses.all_*) can be calculated as the observed value minus the mean of the null distribution, divided by the standard deviation in the null distribution. Calculating the deviation from the null model is commonly performed to express biological differences regardless of the units of the indices (e.g., McCabe et al. 2012; Nakadai and Kawakita, 2016). The value of *P_ses.all_* is close to 0, indicating random deaths of individuals (i.e., ecological drift). In cases in which the value of *P_ses.all_* is greater than 0, the proportion of long-lived individuals (more than three time steps) is larger than that in the null model.

### Case study: the permanent 50-ha forest dynamics plot on BCI in Panama

To demonstrate the use of the new individual-based multiple-unit indices, I analyzed the data from Condit et al. (2019). The permanent 50-ha forest dynamics project plot on BCI in Panama was established in 1981 by Robin Foster and Stephen Hubbell (Hubbell and Foster 1983). Data from this plot have been used in many scientific papers, and descriptions of the area and survey methods are widely available (e.g., Legendre and Condit 2019). This 50-ha plot has been subjected to eight detailed surveys since its inception: in 1982, 1985, 1990, 1995, 2000, 2005, 2010, and 2015. The BCI data were divided into 1,250 quadrats (20 m × 20 m). Harms et al. (2001) classified quadrats into six habitat zones and a zone of mixed habitats (Fig. 3a, Table S1). The six habitat zones are grouped 1–6 and the mixed habitat zone is group 7 (Harms et al. 2001). Detailed information on each group and a map are shown in Figure 3a.

**Figure 3.**
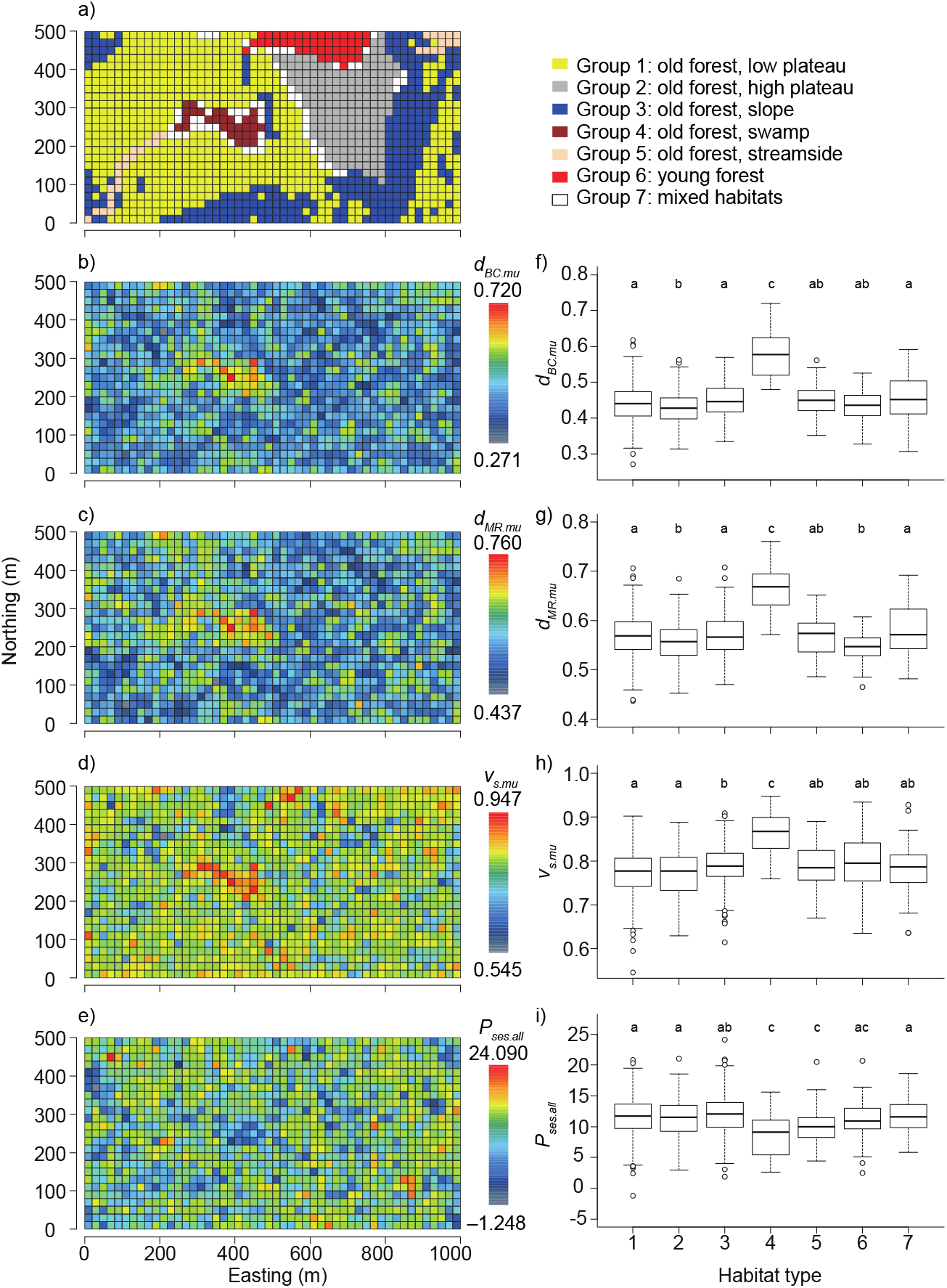
Comparison of the conventional multiple-unit Bray–Curtis dissimilarity (*d_BC.mu_*; b) and f)), individual-based temporal beta-diversity dissimilarity (*d_MR.mu_*; c) and g)), composition variability (*v_s.mu_*; d) and h)), and the degree of deviation in individual persistence from the null model (*P_ses.all_*; e) and i)) among habitat zones a) using the pairwise Wilcoxon rank sum test, which allows for pairwise comparisons between groups with correction for multiple testing. Habitat types labeled with the same letter did not significantly differ in terms of that index based on the Wilcoxon rank sum test with Holm’s method for multiple comparisons. Detailed information about the habitat zones and results of the Wilcoxon rank sum test are shown in Tables S1 and S3, respectively. In a-e), the horizontal and vertical axes indicate easting and northing, respectively.

First, I calculated the values of the two individual-based beta-diversity indices for all possible pairs of time points and then determined their average values 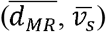 for each quadrat. In addition, I calculated the newly developed multiple-unit indices (*d_MR.mu_, v_s.mu_*) for each quadrat and tested their relationships using simple linear regression to characterize the novel indices. I predicted that the calculated average values would always be lower than the novel multiple-unit indices based on the results of Baselga (2013a), who compared the patterns of the average values of conventional beta-diversity indices and multiple-unit indices. Second, because high individual turnover can result in high compositional variability within a community, to explore how much the values deviate from predictions based on individual turnovers, I tested the relationship between newly developed multiple-unit indices (*d_MR.mu_*) and two previous ones (*d_BC.mu_, d_sor.mu_*). Third, I calculated the degree of deviation in individual persistence from the null model (*P_ses.all_*) based on 100 randomizations via a novel random permutation method (details described above) for each quadrat. I analyzed differences among habitat types in compositional variability (*d_BC.mu_*), the degree of individual turnover (*d_MR.mu_*), compositional variability in individuals undergoing turnover (*v_s.mu_*), and the degree of deviation in individual persistence from the null model (*P_ses.all_*) using the Wilcoxon rank sum test, which allows for pairwise comparisons between groups with correction for multiple testing. The p-values were adjusted by the Holm method. This analysis seeks to afford novel insights on how BCI data can be interpreted using indices based on individual identity information. Finally, because the patterns of individual turnover and persistence may affect community variability, to confirm the effect of individual persistence on community compositional variability, I tested the relationship between the degree of deviation in individual persistence from the null model (*P_ses.all_*) and the conventional multiple-unit dissimilarity index (*d_BC.mu_*) using a generalized additive model fitted by the restricted maximum likelihood (REML) method. I predicted that the conventional multiple-unit dissimilarity index (*d_BC.mu_*) would decrease with increasing deviation of individual persistence from that of the null model (*P_ses.all_*).

All analyses were conducted in R software (ver. 4.0.3; R Development Core Team, 2020). The *betapart* R package (Baselga and Orme 2012) was used to calculate multiple-unit versions of the Bray–Curtis and Sørensen dissimilarity indices. The R package *mgcv* (Wood 2017) was used to fit the generalized additive model to the relationship between the degree of deviation in individual persistence from the null model (*P_ses.all_*) and the conventional multiple-unit dissimilarity index (*d_BC.mu_*). The function *pairwise.wilcox.test* was used to compare the differences among habitats.

## Results

Simple linear regressions showed that *d_MR.mu_* increased significantly with increasing 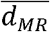 (*adjusted R^2^* = 0.971, *p* < 0.0001; Fig. S1a) and *v_s.mu_* increased significantly with increasing 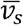 (*adjusted R^2^* = 0.899, *p* < 0.0001; Fig. S1b). Both multiple-unit indices were higher than the average values of the pairwise indices at all times. Both *d_BC.mu_* and *d_sor.mu_* increased significantly with increasing *d_MR.mu_* (*adjusted R^2^* = 0.756, *p* < 0.0001; Fig. S2a and *adjusted R^2^* = 0.444,*p* < 0.0001, respectively; Fig. S2b).

The value of *d_BC.mu_* was significantly higher in group 4 (swamp) than in the other groups (Fig. 3b, f, Table S2a). In addition, no significant difference was found between groups 1 and 3, while the value of group 2 differed significantly from those of groups 1 and 3 (Fig. 3b, f, Table S2a). The *d_MR.mu_* values showed a similar pattern to that of *d_BC.mu_. d_MR.mu_* was significantly higher in group 4 (swamp) than in the other groups (Fig. 3c, g, Table S2b). No clear difference was observed between groups 1 and 3, but the value of group 2 differed significantly from those of groups 1 and 3 (Fig. 3c, g, Table S2b). The *v_s.mu_* value was considerably and significantly higher in group 4 (swamp) than in all other habitats (Fig. 3d, h, Table S2c). In addition, no significant difference in *v_s.mu_* was found between groups 1 (low plateau) and 2 (high plateau), but the value of group 3 (slope) differed from those of groups 1 and 2 (Fig. 3d, h, Table S2c). There were no significant differences in *P_ses.all_*, among groups 4 (swamp), 5 (streamside), and 6 (young forest), but the values were significantly lower in groups 4 and 5 than in the other groups (i.e., groups 1, 2, 3, and 7) (Fig. 3e, i, Table S2d). Moreover, no significant difference was found between groups 1 (low plateau) and 2 (high plateau), but group 3 (slope) differed significantly from groups 1 and 2 in terms of *P_ses.all_* (Fig. 3e, i, Table S2d).

In the generalized additive model, compositional variability (*d_BC.mu_*) showed a significant response to the degree of deviation in individual persistence from the null model (*P_ses.all_*: edf = 3.951, *F* = 31.59, *adjusted R^2^* = 0.112, *p* < 0.0001), with the values of the compositional variability index constant from *P_ses.all_* of approximately 12 to 25 and tending to increase from *P_ses.all_* of approximately 12 to 0 (Fig. 4).

**Figure 4.**
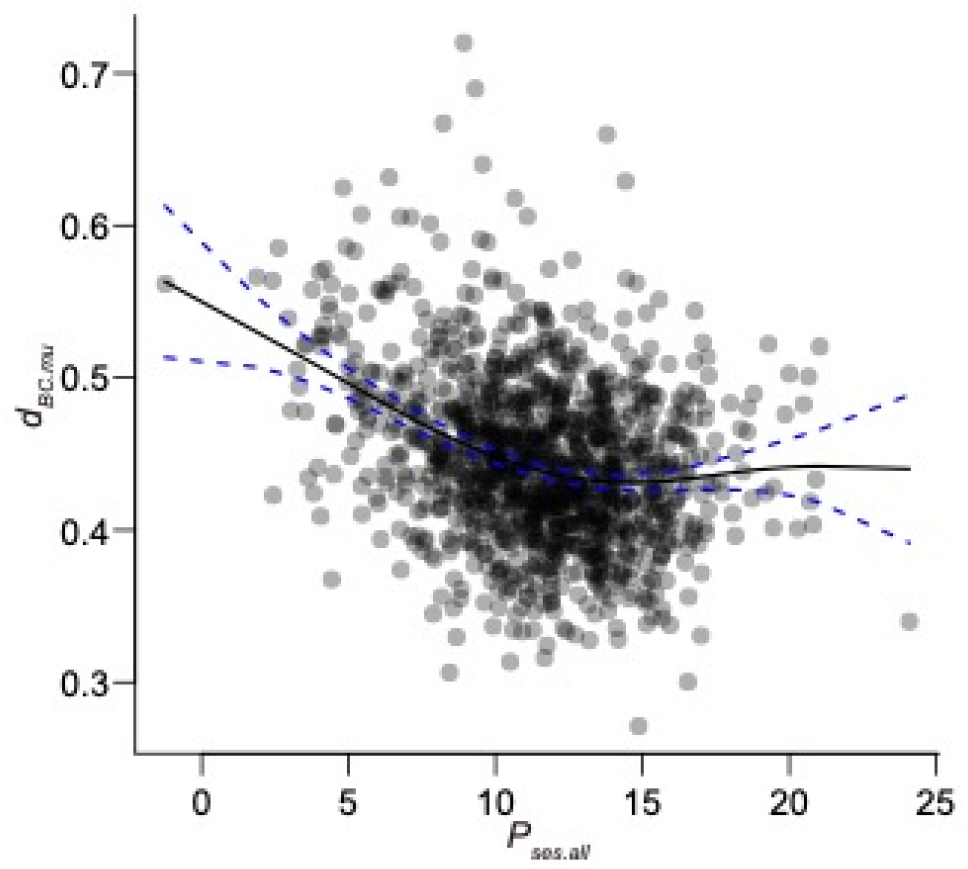
The relationship between the degree of deviation in individual persistence from the null model (*P_ses.all_*) and the conventional multiple-unit dissimilarity index (*d_BC.mu_*). The solid black line was drawn using a generalized additive model fitted using the restricted maximum likelihood method (*adjusted R^2^* = 0.112, *p* < 0.0001, *edf* = 3.951, *F* = 31.59). The blue dashed lines represent the 95% confidence intervals.

## Discussion

Individual turnover is a major driver of compositional change and variability within a community, although it has rarely been discussed. Higher individual turnover clearly facilitates higher variability in community composition; thus, compositional variability over time can occur even when considering only ecological drift. Hubbell’s unified neutral theory of biodiversity (Hubbell 2001) postulates ecological neutrality among species within communities and emphasizes the role of ecological drift in shaping biodiversity patterns. To elucidate compositional variability, the impact of individual turnover (i.e., ecological drift) should be excluded. However, almost no methods exist to quantify the impact of individual turnover on compositional variability. The individual-based indices introduced herein can assess compositional variability considering the contribution of individual turnover. In addition, information about individual identity across multiple time steps provides opportunities to test individual persistence. Within and among species, some individuals live longer (i.e., individuals persist in the community for a longer period) than others. The patterns of individual persistence can be tested for their degree of deviation from expected patterns due to stochastic drift using the novel null model introduced here. This model can add a new assessment axis to empirical studies of community dynamics.

In the present study, I applied both conventional and novel indices, as well as the degree of deviation based on the null model to the BCI plot. The average values of both 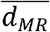 and 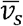 across all pairs are always lower than multiple-unit indices (i.e., *d_MR.mu_* and *v_s.mu_*; Fig. S1), in accordance with the findings of Baselga (2013a). The results of the case study clearly demonstrated that the novel individual-based index was significantly correlated with conventional incidence- and abundance-based indices (Fig. S2); however, variations among communities still exist. The value of *v_s.mu_*, which represents compositional variability after considering individual turnover, differed among habitat zones, especially for group 4 (swamp), which differed significantly from all other habitats (Fig. 3a, c, Table S1). Both the degree of compositional variability (*d_BC.mu_*) and individual turnover (*d_MR.mu_*) were significantly higher in group 4 than in all other habitats. There was significant compositional variability in the swamp even after controlling for individual turnover. Thus, individual turnover is a major driver, but a large component of variability remains unexplained by individual turnover. The environmental stability of each plot is a possible source of the unexplained effect. For example, Nakadai (2020) identified compositional changes after consideration of the individual turnover that was clearly associated with the degree of climate change (i.e., rate of temperature change) across the Japanese Archipelago. Compositional change over time in this swamp habitat has also been reported in terms of responses to a drought event at the BCI plot (Legendre and Condit 2019). In addition, although *d_BC.mu_* and *d_MR.mu_* were significantly higher in groups 1 (low plateau) and 3 (slope) than in group 2 (high plateau), *v_s.mu_* did not follow the same pattern. Instead, *v_s.mu_* was significantly lower in groups 1 (low plateau) and 2 (high plateau) than in group 3 (slope). These results suggest that the high plateau (group 2) has lower individual turnover, probably because it is isolated and stable and consequently has lower compositional variability. However, when considering individual turnover, the slope (group 3) differs from both the low and high plateau (groups 1 and 2).

The degree of deviation in individual persistence was lower in group 4 (swamp) than in groups 1, 2, 3, and 7, but not group 5 (streamside) or 6 (young forest), suggesting that the three habitat types (i.e., swamp, streamside, and young forest) may be approaching their expected mortal patterns due to ecological drift. Moreover, based on the generalized additive model, compositional variability was associated with the degree of deviation in individual persistence from the null model (Fig. 4). This suggests that an increase in the proportion of the long-lived individuals (i.e., higher values of *P_ses.all_*) reduces compositional variability, even if the level of individual turnover remains the same. Both the results and the interpretation thereof indicate that the novel approach introduced in the present study affords new insights into the forest dynamics of community composition. In previous studies, individual persistence was recognized in community dynamics as a factor facilitating stable coexistence among species and buffering population growth (i.e., the storage effect) (Chesson and Warner 1981; Chesson 2000). The effect of buffered population growth is expected to be low compositional variability and turnover of individuals over time throughout the community. This prediction is consistent with the results related to *P_ses.all_*. In other words, habitat types with higher *P_ses.all_* (except groups 4, 5, and 6) were better stabilized by the storage effect.

The novel approaches introduced in the present study focus on datasets collected via individual tracking. I chose one of the most representative and widely employed datasets (the BCI forest plot; Hubbell and Foster 1983) as an example. It is difficult to apply the new methods to taxa in which individuals are not identified (e.g., aquatic plankton communities) unless they are all experimentally tracked. However, I emphasize that my approach can be applied in many other scenarios. For example, the Forest Global Earth Observatory (ForestGEO; https://forestgeo.si.edu/) is a global network of scientists and forest research sites including the BCI plot; data from 72 sites across 27 countries are available (Davis et al. 2021). On the regional scale, the Monitoring Sites 1,000 Project launched by the Ministry of the Environment, Japan (Ishihara et al. 2011) includes 60 forest plots across the Japanese archipelago (Ishihara et al., 2011; Nakadai 2020). Both the ForestGEO and the monitoring 1000 projects feature individual tracking; my methods are applicable. In addition, recent developments in remote sensing techniques have rendered it possible to identify individual trees (e.g., Guillén-Escribà et al. 2021; Weinstein et al. 2021), although not (yet) to track individuals. In the near future, remote sensing will yield individual tracking datasets on a global scale. It is sometimes difficult to identify clonal plant species or individuals with multiple stems. However, if the target shifts from the individual to the stem, my methods are applicable. Therefore, depending on the purpose and status of a study, it may be possible to change the analytical resolution and the target. Finally, the concept of individual-based beta diversity is applicable not only to plant community datasets but also to animal datasets (Nakadai 2020). Specifically, datasets on bird ringing, other classical mark–recapture procedures, and biologging, which record animal ranges and movements, can be analyzed using my methods because multiple time-series datasets collected from the same place can track individual turnover across time and space (Nakadai 2020).

Although temporal beta diversity has attracted much attention, especially at large spatial scales (Gotelli et al. 2017; Brice et al. 2019; Magurran et al. 2019), it is simply an extension of spatial beta diversity. Both methodological and conceptual frameworks rooted in community dynamics should be used. Above all, information from “individuals” has been absent from empirical studies of temporal dynamics in community ecology and biodiversity. On the other hand, recent advances in modern coexistence theory (Chesson 2000) suggest that the impact of individuals across generations (e.g., individual persistence of adult trees) is a storage effect that facilitates species coexistence and stabilizes community dynamics. In this context, the methods proposed here are expected to be useful for addressing a wide range of research questions related to temporal changes in biodiversity, especially studies using forest monitoring data to track individuals, by bridging gaps between recent theoretical advances and accumulated empirical data.

## Acknowledgments

Financial support was provided by the Japan Society for the Promotion of Science (grant 18J00093).

## Declarations

### Data Availability

Data are available from the Dryad Digital Repository (https://doi.org/10.15146/5xcp-0d46; accessed on 10^th^ of April in 2020) (Condit et al. 2019).

## Author contributions

R.N. conceived the study, designed the methods described herein, analyzed the empirical data, and wrote the manuscript.

## Appendices

**Figure S1** Correlations between novel multiple-unit indices and the average values of conventional pairwise indices (a) *d_MR_-d_MR.mu_; adjusted R^2^* = 0.971, *p* < 0.0001, b) *v_s_-v_s.mu_; adjusted R^2^* = 0.899, *p* < 0.0001).

**Figure S2** Correlations between a novel individual-based multiple-unit index (*d_MR.mu_*) and conventional multiple-unit indices (*d_BC.mu_*) (a) *d_BC.mu_-d_MR.mu_; adjusted R^2^* = 0.756, *p* < 0.0001, b) *d_sor.mu_-d_MR.mu_; adjusted R^2^* = 0.444, *p* < 0.0001).

**Figure S3** Examples of patterns of individual persistence with the corresponding values of *P_ses.all_*. The x-axis indicates the number of time periods of individual persistence. The y-axis indicates the number of individuals persisting. a) The case of Quadrat No. 1221 (*P_ses.all_* = 2.403), b) the case of Quadrat No. 0322 (*P_ses.all_* = 24.090).

**Table S1** Habitat clarification in Harms et al. (2001). Group 1: old forest, low plateau; group 2: old forest, high plateau; group 3: old forest, slope; group 4: old forest, swamp; group 5: old forest, streamside; group 6: young forest; and group 7: mixed habitats.

**Table S2** Comparisons of the conventional multiple-unit Bray–Curtis dissimilarity (*d_BC.mu_*: a)), the individual-based temporal beta-diversity dissimilarity index (*d_MR.mu_*: b)), composition variability (*v_s.mu_*: c)), and the degree of deviation in individual persistence from the null model (*P_ses.all_*: d)) among habitat zones using the pairwise Wilcoxon rank sum test, which allows for pairwise comparisons between groups with correction for multiple testing. The tables correspond to f-i) in Figure 3.

**Supplementary File 1** R script for the individual-based multiple-unit beta-diversity calculations performed using sample data.

