## Supplementary figures and images for "Individual-based multiple-unit dissimilarity: novel indices and null model for assessing temporal variability in community composition"

### FIgure S1

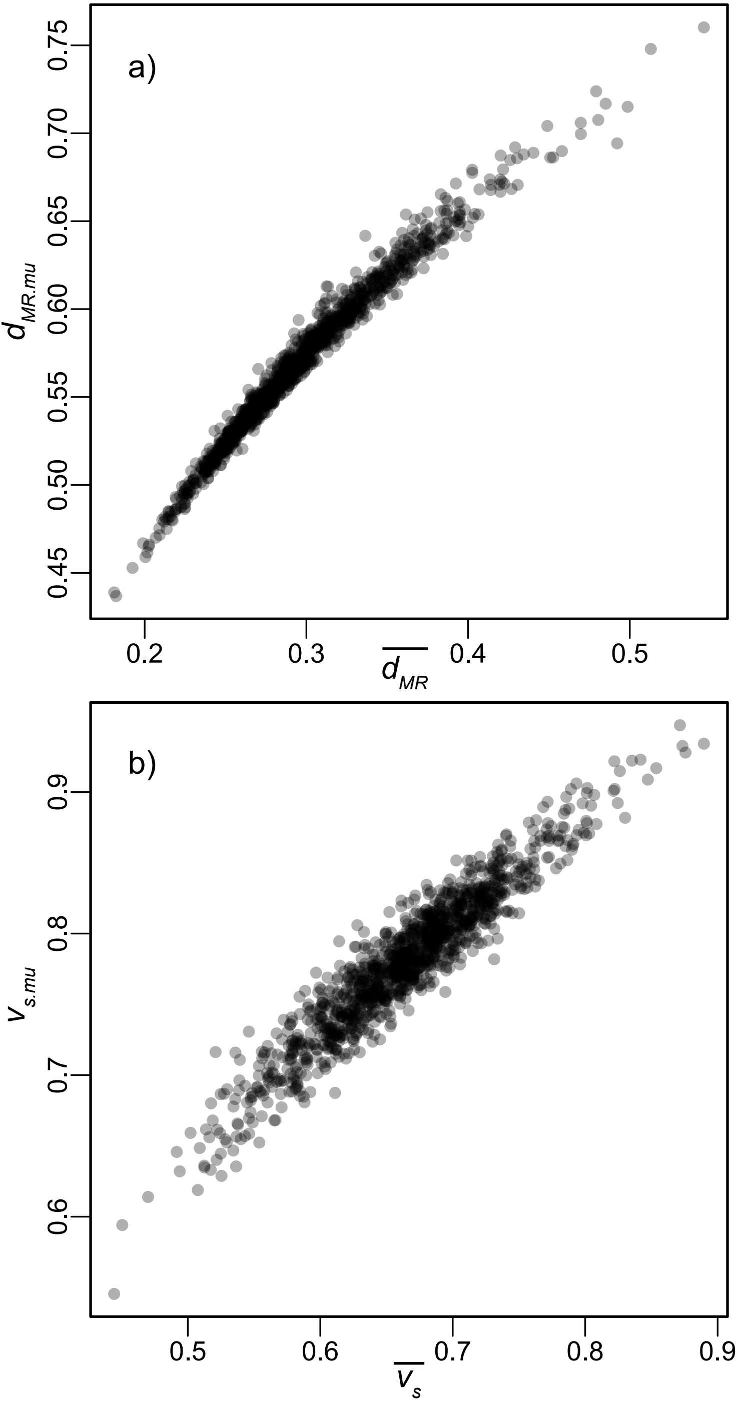

### Figure S2

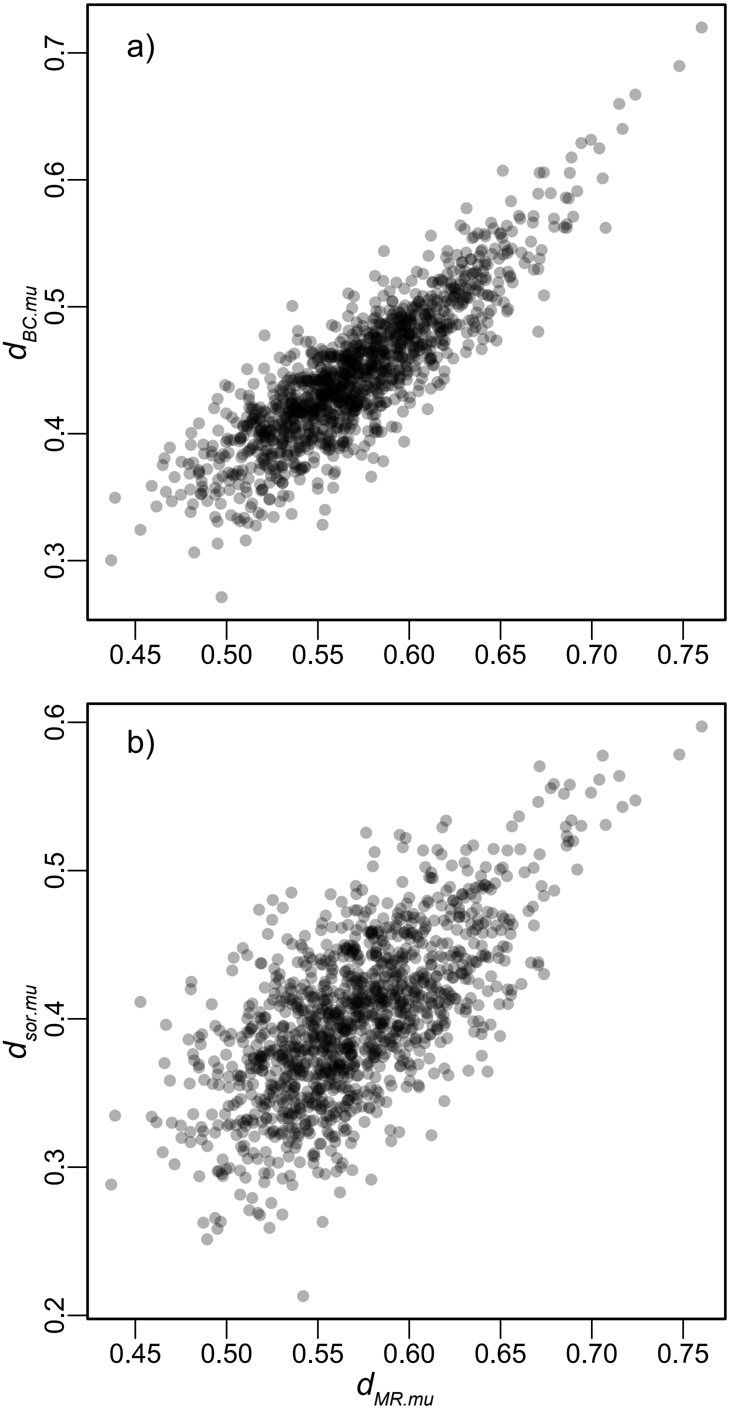

### Figure S3

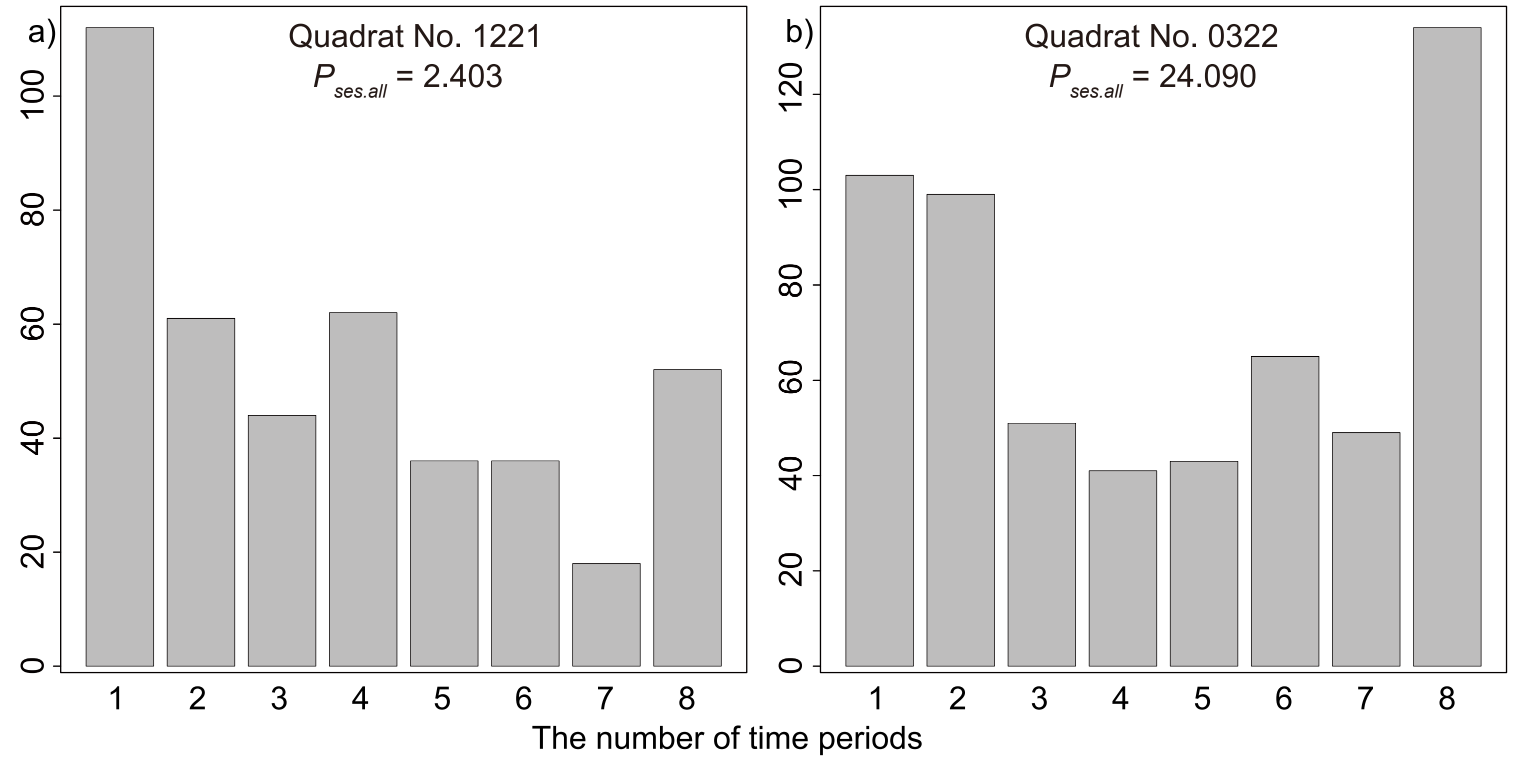
